# Transposable elements identify previously overlooked regions undergoing parallel evolution in response to diverse selective pressures

**DOI:** 10.1101/2024.04.06.588382

**Authors:** Sara Guirao-Rico, Josefa González

## Abstract

Evolve and Resequence (E&R) studies have contributed to closing the gap between adaptive phenotypic traits and their underlying genomic basis. Unlike other popular approaches, E&R studies provide a unique opportunity to understand the degree of repeatability and constraints of the evolutionary process. Still, one of the major challenges is to accurately identify the genetic variants targeted by selection. Traditionally, E&R studies have only explored part of the variation present in the genomes, SNPs, thus neglecting other sources of adaptive variation such as transposable elements (TEs). In this work, we analyzed the Pool-seq data of five published Drosophila melanogaster E&R studies in the context of TEs. We found some candidate TEs inserted in the same or nearby regions to significant SNPs/genes. As such, these TEs should be added to the set of possible targets of selection. However, we also found a subset of significant TEs that were not associated with previously identified significant SNPs/genes. Most of these TEs are inserted within or flanked by genes related to the trait of interest and thus they are the only potential targets of selection identified in these regions. Functional enrichment analysis of the genes located nearby all candidate adaptive TEs allowed to identify new relevant biological functions. Finally, the replicability of candidate TEs was similar or higher than that reported for SNPs. We argue that TE variants should also be considered as candidate targets of selection in E&R studies and we also provide guidelines to improve the detection of TEs as targets of selection.

## INTRODUCTION

The link between genotype and phenotype is still an elusive step in Evolutionary Biology studies. Over the years, several approaches, both bottom-up and top-down, have tried to identify and quantify the fraction of genetic variation enabling adaptation in natural populations mostly in a retrospective way (Savolainen, et al. 2013). Experimental Evolution (EE) provides a very valuable research framework to study the mechanisms of evolution through a prospective approach (Bailey, et al. 2016). EE studies allow us to test theoretical predictions about which are the traits or phenotypes relevant for adaptation to particular factors in a specific situation where some aspects that are impossible or difficult to handle in natural populations are controlled *e.g.* simulating an ancestral environment or keeping a specific temperature constant through time. Thus, EE studies allow us to observe the evolution of a particular trait in a specific environmental context *e.g*. establishing the link between body size and environmental temperature. Advances in high-throughput methods, *e.g*. genome sequencing and mapping, have made possible to apply genomics to EE, the so called, Evolve and Resequence studies (E&R), allowing the identification of genetic variants or at least narrowing the genomic regions underlying the traits of interest and hence, connecting the genotype with the phenotype (Schlötterer, et al. 2015; Long, et al. 2015). In addition, E&R studies allow us to assess the patterns of parallelism in the response to selection and thus to evaluate how predictable adaptation is (Schlotterer 2023). As such, E&R studies are one of the experimental approaches used to assess climate change adaptation (Hoffmann, et al. 2023).

Some of the requirements to perform E&R studies, such as short generation times, a well-annotated genome, and the availability of genetic tools for functional studies to eventually link genotypic to phenotypic variation, imposed some limits on the range of organisms that have been explored. Many of the E&R studies to date have been performed on the model organism *Drosophila melanogaster*. Several traits under selection have been studied using this species *e.g.* starvation and desiccation resistance, diet and food consumption, lifespan and late age fertility, egg size, immunity, courtship song, body size, hypoxia, darkness and accelerated development (Table S1). While Drosophila continues to be a premier system for E&R studies, the range of organisms in which this experimental approach has been successfully applied continues to increase and includes diverse organisms, such as bacteria, yeast, plants and nematodes, among others (*e.g.* Burghardt, et al. 2018; Phillips, et al. 2020; Johnson, et al. 2023; Eoche-Bosy, et al. 2017).

It is well documented that transposable elements (TEs) are an important source of genetic variation and that in many cases TEs are involved in rapid adaptive responses to challenging environmental conditions (Casacuberta and González 2013; Stapley, et al. 2015; Quadrana, et al. 2019; Baduel and Quadrana 2021; Thieme, et al. 2022). In addition, there is increasing evidence for the functional impact of individual TEs inserted near genes involved in stress response (Makarevitch, et al. 2015; Horváth, et al. 2017; Rech, et al. 2019; Villanueva-Cañas, et al. 2019; Ullastres, et al. 2021; Merenciano and Gonzalez 2023). Despite its adaptive potential, and as in other genomic studies, TEs have been neglected in E&R studies, mainly due to technological limitations associated with their repetitive nature. To date, and to the best of our knowledge, only two studies have considered other sources of variation different from SNPs in the response of *D. melanogaster* to shock temperature, starvation and desiccation (Michalak, et al. 2019) and longevity (Fabian, et al. 2021). However, these studies focused on the potential adaptive role of TEs from a quantitative perspective, mainly testing whether the TE load and the number of TEs per family significantly differed between control and selected populations.

In this work, we analyzed the data of five published E&R studies evolved populations under four different selective regimes, increased lifespan and late-age fertility (Remolina, et al. 2012), dietary selection (Reed, et al. 2014), desiccation resistance (Griffin, et al. 2017, Kang, et al. 2016), and starvation resistance (Hardy, et al. 2018), in the context of TEs (Table 1). We selected, among the several published studies, those with relatively high sequencing coverage, number of individuals and replicates, and pair-end read sequencing, for statistical power reasons (Table 1 and Table S1). We have explored TEs in these five E&R studies to search for hallmarks of positive selection associated with both reference and non-reference TE insertions, and to assess to what extent the same TEs are selected for the trait of interest repeatedly across replicates.

**Table 1.** Summary of the main features of each E&R experiment analyzed in this work.

| **Study** | **Trait/Stress** | **Number of individuals in the starting population** | **Generations of selection** | **Pools sequenced (replicates)** | **Number of individuals per pool** | **Coverage per pool** |
| --- | --- | --- | --- | --- | --- | --- |
| **Remolina** | Lifespan and late-age fertility | 400 | 50 | 3 independent selection-control pairs of populations | 100 | 40-50x |
| **Reed** | Dietary selection | 200 | 14-17 | 7 baseline pools (normal diet); 2 replicates of populations selected for each diet: high fat, high sugar, and low sugar. | baseline: 288; pool: 144-180 | 150-463x |
| **Griffin** | Desiccation resistance | 60 | 13 | 1 mass bred; 5 control and 5 selection populations | mass bred: 200, pool: 50 | mass bred: 130x, pool: 22x-72x |
| **Kang** | Desiccation resistance | 120 | 48 | 3 control and 3 selection populations | 100 | 31-38x |
| **Hardy** | Starvation resistance | 400 | 83 | 3 control and 3 selection populations | 100 | 5.9-72x |

## RESULTS

We analyzed the Pool-seq data of five published E&R studies corresponding to evolved populations under four different selective regimes (increased lifespan and late-age fertility, dietary selection, desiccation resistance, and starvation resistance), in the context of TEs (Table 1). Besides the trait of interest, the evolved populations also differed in the number of replicates, number of individuals in the starting population and in the sequenced replicates, number of generations under selection and other aspects of the experimental design that can be relevant to assess the replicability of the adaptive response and the power to detect and identify the causal selected mutations (Table 1). We analyzed both reference TE insertions, *i.e.* TEs that have been annotated in the *D. melanogaster* reference genome, and non-reference TE insertions, *i.e.* TEs present in the strains analyzed in this work but not annotated in the reference genome. To detect presence/absence and estimate frequencies of reference TE insertions, we used Tlex-3 (Bogaerts-Márquez, et al. 2020), while to identify non-reference TE insertions we used two different software TIDAL and TEMP (Rahman, et al. 2015; Zhuang, et al. 2014). Because analyzing TEs that are detected by two software increases the TE detection sensitivity (Vendrell-Mir, et al. 2019), we considered non-reference TEs that were detected both by TEMP and TIDAL and the reference TEs detected by T-lex3: the TEMP&TIDAL+T-lex3 dataset. However, considering only the non-reference TEs detected by two software can also introduce biases, as individual software can identify TEs that are not detected by other methods (Nelson, et al. 2017). We thus also analyzed a dataset including the non-reference TEs identified by TEMP and the reference TEs identified by T-lex3: the TEMP+T-lex3 dataset. We chose TEMP as this software not only detects presence/absence of TEs in pools but also estimates TE frequencies. In general, the number of non-reference TEs detected by both software were around half or a third of the values of those found by each software independently, with TIDAL detecting a higher number of TEs compared with TEMP in most of the studies (Table 2A and Table S2). Note that for the Reed dataset, T-lex3 was not able to detect presence/absence of reference TE insertions and TEMP detected a much smaller number compared with TIDAL (see Material and Methods), and as a result, the number of TEs analyzed for this dataset is much smaller compared with the other datasets (Table 2A). For the other four studies, the total number of TEs detected per dataset and pool ranged between 899-1713 for TEMP&TIDAL+T-lex3 and 1287-3022 for TEMP+Tlex3.

**Table 2.** Number of detected and significant TEs and their change in frequencies. A) Average number of non-reference TEs, detected by TEMP, TIDAL, and both software, and reference TEs detected by T-lex3 in all the pools analyzed. B) Average number of TEs detected in control (or MB: mass bred or BS: baseline) and selected pools by TEMP&TIDAL+Tlex3. TEs detected by TEMP+Tlex-3 are given in parenthesis. C) Average number of TEs detected by TEMP&TIDAL+Tlex3 with significant frequency changes. TEs detected by TEMP+Tlex-3 are given in parenthesis. Note that each replicate is analyzed independently. D) Number of significant TEs detected by TEMP&TIDAL+Tlex3 that significantly increased in frequency in >1, only 1 replicate or private. TEs detected by TEMP+Tlex-3 are given in parenthesis. E) Average difference in frequency between control and selected pools (or MB or BS). TEs detected by TEMP+Tlex-3 are given in parenthesis.

|  |  | Remolina | Reed | Griffin | Kang | Hardy |
| --- | --- | --- | --- | --- | --- | --- |
| **A** | **Average number of non-reference TEs detected by TEMP** | 1446.33 | 225.38 | 1497.55 | 1064.40 | 1315.67 |
|  | **Average number of non-reference TEs detected by TIDAL** | 1277.67 | 2967.85 | 1664.55 | 1141.00 | 2368.83 |
|  | **Average number of non-reference TEs detected by TIDAL and TEMP** | 710.50 | 82.46 | 537.91 | 608.17 | 801.20 |
|  | **Average number of reference TEs detected by Tlex** | 551.33 | NA | 630.91 | 586.33 | 679.50 |
| **B** | **Average number of TEs in control pools (or MB or BS)** | 1277.33<br>(1973.33) | 31.71<br>(79.71) | 1067.17<br>(2063.33) | 1281.33<br>(1829.33) | 1578.33<br>(2119) |
|  | **Average number of TEs in selected pools** | 1246.33<br>(2022) | 141.67<br>(395.33) | 1290.8 (2206.6) | 1107.67<br>(1528.33) | 1427<br>(1871.33) |
| **C** | **Average number of significant TEs** | 47.67 (76) | 1.17 (1.5) | 12.4 (18.8) | 40.89 (61) | 95.33 (123.33) |
|  | **Average number of significant TEs with increase in frequency** | 15.67 (24.33) | 0.5 (0.67) | 10.8 (15.8) | 34.67 (51.67) | 76.89 (109) |
|  | **Average number of significant TEs with decrease in frequency** | 32 (51.67) | 0.67 (0.83) | 1.6 (3) | 6.22 (9.33) | 18.44 (21) |
| **D** | **Number of significant TEs with increase in frequency in >1 replicate** | 0 (1) | 0 (0) | 10 (12) | 58 (80) | 85 (121) |
|  | **Number of significant TEs with increase in frequency in one replicate** | 6 (6) | 3 (4) | 18 (36) | 17 (54) | 141 (186) |
|  | **Number of significant TEs with increase in frequency and private of one replicate** | 6 (6) | 0 (0) | 8 (16) | 2 (14) | 83 (103) |
| **E** | **Average difference in frequency between control (or MB and BS) and selected pools** | 0.53 (0.49) | 0.50 (0.50) | 0.31 (0.32) | 0.66 (0.64) | 0.46 (0.56) |

### TE abundance does not significantly change between control and selected pools

Quantifying if there are differences in TE abundance between control (or mass bred or baseline) and selection pools can shed light on the evolutionary forces to which these genetic elements are subject and whether the change in TEs abundance is trait-dependent. In previous studies, TEs abundance was reported to increase in desiccation resistance compared to control conditions (Michalak, et al. 2019), and in long-lived populations compared with non-selected controls (Fabian, et al. 2021), while it decreased in heat-shock resistance compared with control conditions (Michalak, et al. 2019). Across the five studies and the two TE datasets analyzed here, we found that only the increase in TEs abundance in response to dietary selection was significant when the TE dataset identified by TEMP+T-lex3 was analyzed (Wilcoxon test, two-sided, *P*-value = 0.005; Figure 1, Table 2B and Table S2). Thus, overall, we did not find significant differences in the abundance of TEs in response to the selected traits analyzed.

**Figure 1.**
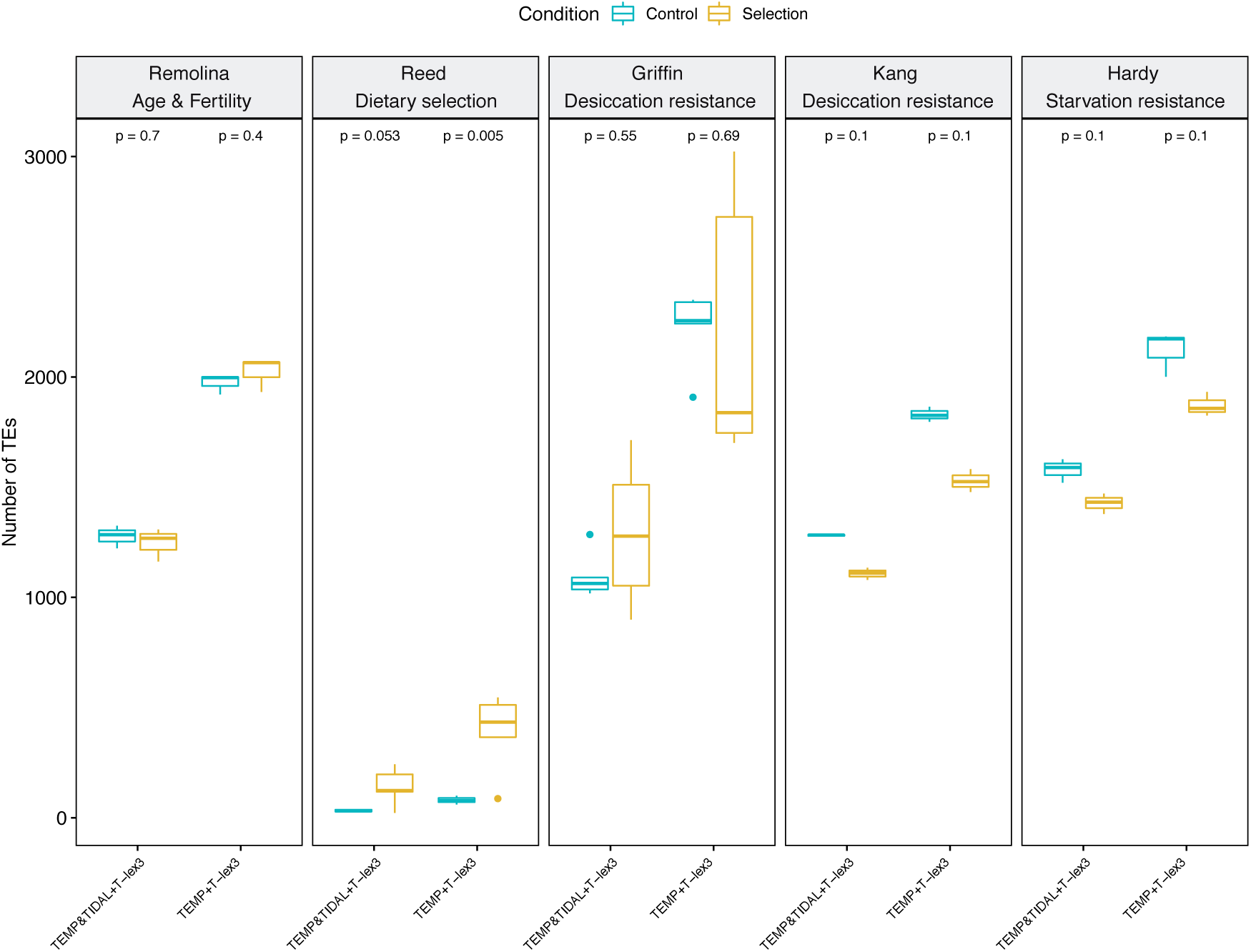
Differences in the number of TEs between the control and selected conditions for each E&R study analyzed and their associated *P*-values.

### The majority of TEs with a significant frequency change between control and selected pools increased in frequency in response to selection

To assess the direction of selection acting on TEs, we calculated the change in TE frequencies between control and selected pools and its direction considering each replicate independently. As mentioned above, the number of TEs analyzed for the Reed dataset was small, and on average only one of the TEs analyzed changed significantly in frequency between control and selection pools (Table 2C). For the other four datasets, the average number of TEs that significantly change their frequency between control and selection pools after correcting by Bonferroni (*P*-value < 0.05) ranged from 12 to 95 for the TEMP&TIDAL+T-lex3 dataset, and from 19 to 123 for the TEMP+T-lex3 dataset (Table 2C and Table S3). When we considered the number of significant TEs according to the direction of the frequency change between control and selected pool replicates, in three out of the four studies we found a higher number of TEs whose frequencies increased from control to selection pools. Note that the two studies with a similar experimental design, Hardy and Kang, showed the highest number of TEs with significant increase in their frequencies, suggesting that the experimental design might affect the power to detect significant changes (Table 2C and Table S3). These results might suggest that positive selection is more prevalent than negative selection for the TEs analyzed in this work that significantly change in frequency.

### Replicability of candidate TEs is similar or higher than replicability of candidate SNPs in the majority of studies

We identify candidate TEs as those that significantly increase in frequency between control and selected pools consistently across replicates or in a single replicate. The number of candidate TEs varied across studies and ranged from 3 to 226 (4 and 307 when considering TEM+T-lex3 dataset) (Table 2D). These candidate TEs were distributed across all major chromosomal arms (Figure 2).

**Figure 2.**
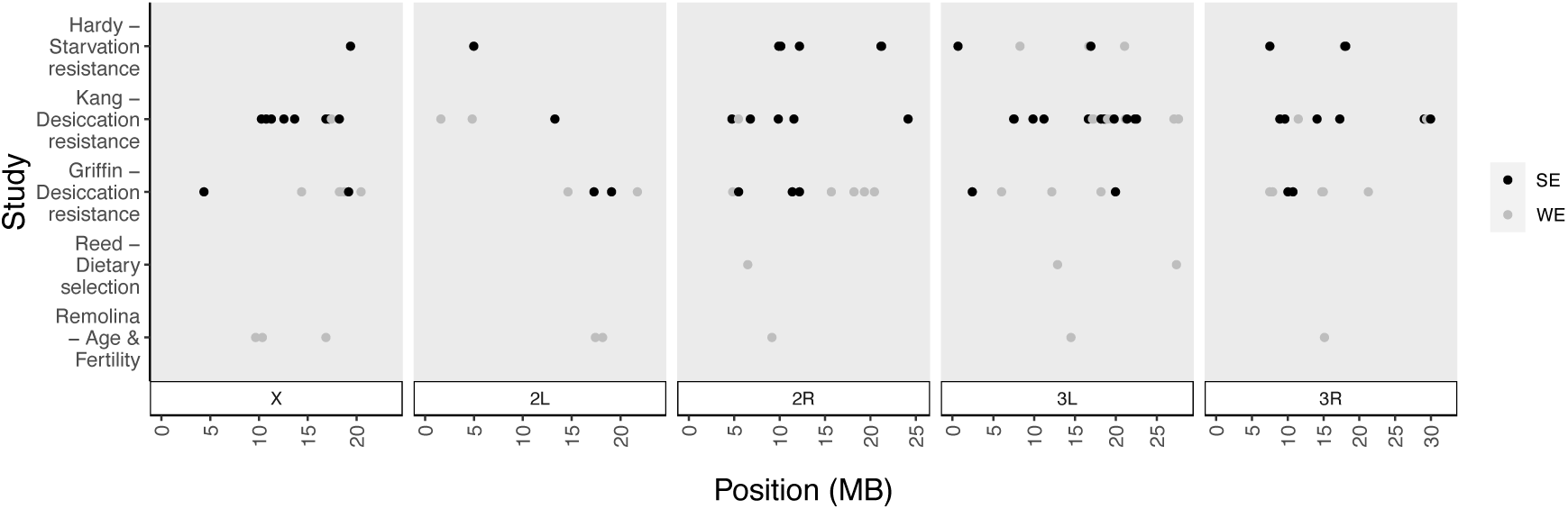
Genomic distribution of the significant TEs identified in each of the five studies analyzed. TEs are classified according to their level of supporting evidence for each E&R study in strong evidence (*SE*) and weak evidence (*WE*).

As mentioned above, E&R studies allow identifying patterns of parallel evolution. In the case of SNPs, three of the studies analyzed in this work quantified the percentage of selected SNPs present in more than one replicate: 9% in Griffin, 36% in Reed and 17.7% in Hardy. While the other two studies, Remolina and Kang, reported high levels of replicability in the SNPs under selection. As one of the goals of this work is to assess if TEs show patterns of parallel selection across replicates, we thus searched for TEs whose frequency significantly and consistently increased from control to selected pools in more than one control-selection replicate. Three studies, Griffin, Kang and Hardy showed TEs with a significant increase in frequency in more than one replica (36%, 77% and 38% respectively for TEs detected with TEMP&TIDAL+T-lex3 and 25%, 60% and 39% for TEs detected with TEMP+T-lex3) (Figure 3 and Tables 2D and S4C-E). Note that these levels of replicability are similar or higher than those reported for SNPs. On the other hand, in Remolina and Reed all the significant TEs were found in only one replicate (Figure 3 and Tables 2D, S4A and S4B). In the case of the Reed study, the lack of replicability is most probably due to the low number of TEs detected (Table 2A). Again, and in contrast with the others, the two studies with the same experimental design (Kang and Hardy) found a substantial number of significant TEs shared among replicates (58-80 and 85-121, respectively) (Figure 3 and Tables 2D, S4D-E). Note that, Kang and Hardy are indeed the two studies with the higher number of significant TEs compared with the other E&R experiments analyzed here (Table 2D and Tables S4D-E), which is similar to what has been found when analyzing SNPs.

**Figure 3.**
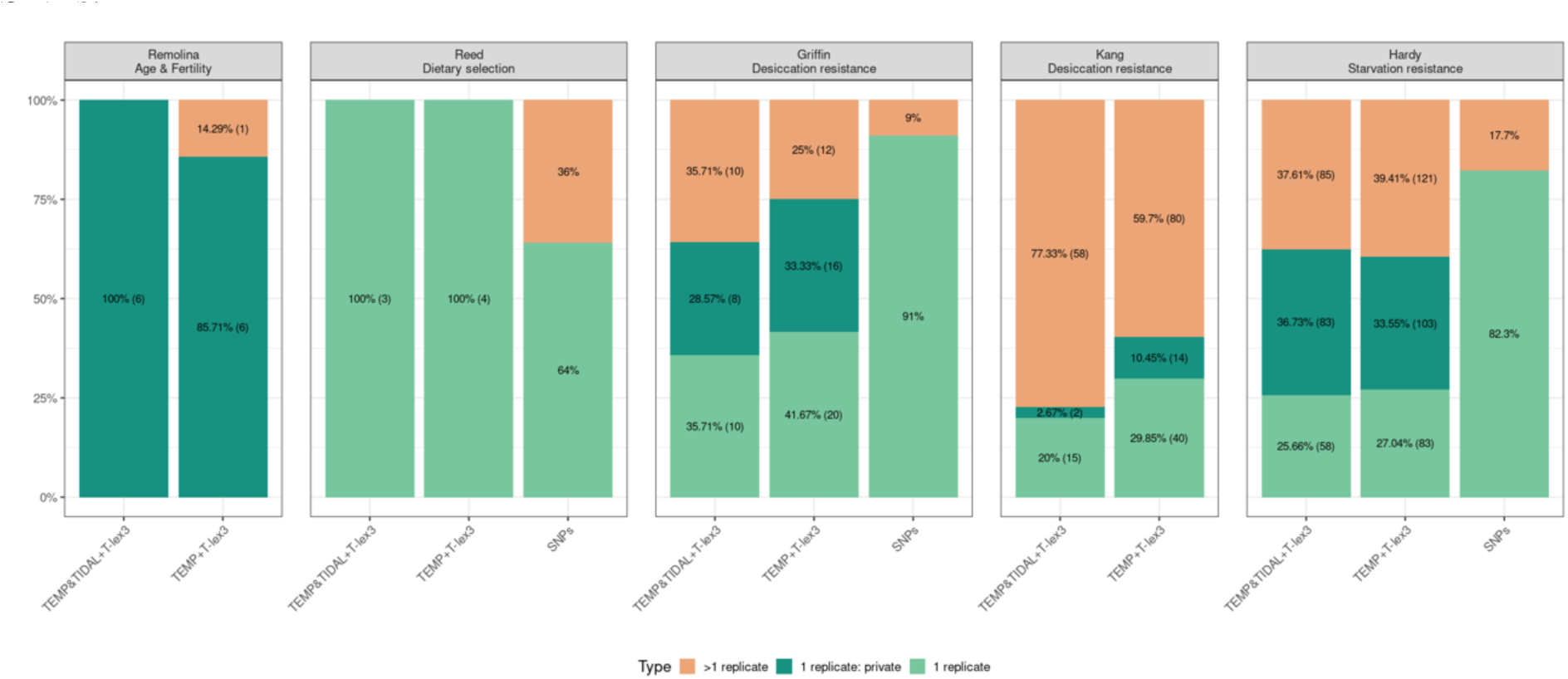
Proportion of significant TEs found in more than one replicate, one replicate and private for each E&R study. When available, the proportions are also given for SNPs.

While focusing only on TEs that are significant in more than one replica helps identify TEs with strong evidence of selection, exploring only parallel patterns of selection could lead to ignoring genomic variation that could be contributing to the traits under selection (Griffin, et al. 2016). This is more relevant in the case of TEs, because most of them are present at very low population frequencies (*e.g.* Rech, et al. 2022), which can lead to some of them not even being present in some replicates due to the stochasticity in the sampling of individuals from the starting population. Thus, besides TEs that were present in several replicates but were only significant in one, we also identified TEs that were significant in one replicate but were not present in the control pools of the other replicates (hereafter private TEs). In Remolina, all the significant TEs increasing in frequency in one replicate were indeed private (Figure 3 and Table 2D; see Material and Methods). In Griffin and Hardy, the percentage of TEs increasing in frequency in only one replica that were private was also considerable (44.4% and 58,9%, respectively Figure 3 and Table 2D). Note that in Hardy, and similarly to the results found with SNPs, the high number of significant TEs that were private of one selection pool was mainly due to a high number of significant TEs in selection pool A (Tables S4E7-E9, Hardy, et al. 2018). Thus, a substantial proportion of the TEs that increased in frequency in only one replicate were private, with Hardy showing the higher proportion of significant private TEs (58.9% for the TEMP&TIDAL+T-lex3 dataset and 55.4% for the TEMP+T-lex3 dataset).

Overall, the replicability of TEs is similar or higher than the replicability of SNPs in three of the five studies analyzed (Figure 3 and Table 2D). In Reed, the lack of replicability observed for TEs is probably due to the low number of TEs analyzed while in Remolina, we found that TEs increasing in frequency in only one replicate were indeed present only in the control pools of that replicate, which explains the lack of replicability.

### Average increase in frequency of candidate TEs is similar or higher than that of candidate SNPs

The average change in TE frequencies between control and selected pools ranged from 0.31 to 0.66 for the TEMP&TIDAL+T-lex3 dataset and from 0.32 to 0.64 for the TEMP+T-lex3 dataset (Figure 4 and Table 2E). In the case of the Remolina study (Remolina, et al. 2012), the values were above those reported using SNPs, but in line with those reported in other E&R studies (Reed, et al. 2014; Griffin, et al. 2017). Similarly, in Reed, the observed change in frequency of candidate TEs is far above the required change in frequency of 0.3 to consider a SNP as a candidate, although this average was obtained only from a small number of candidate TEs (3-4 candidates; Table 2D). In contrast, the average change in frequency in the case of the Griffin study was below the median frequency change per replicate in candidate SNPs (0.31-0.32 for TEs vs 0.48-0.62 for SNPs). Finally, although Kang and Hardy do not provide the change in frequency of the candidate SNPs, we found that the average change in frequency of candidate TEs in these two studies was substantial, and similar to the other E&R studies analyzed in this work (Table 2E). This relatively high change in frequency is still high even if we only considered those significant TEs found in the three replicates (Kang: 0.62 for TEMP&TIDAL+T-lex3 and 0.64 for TEMP+T-lex3; Hardy: 0.47 for TEMP&TIDAL+T-lex3 and 0.56 for TEMP+T-lex3).

**Figure 4.**
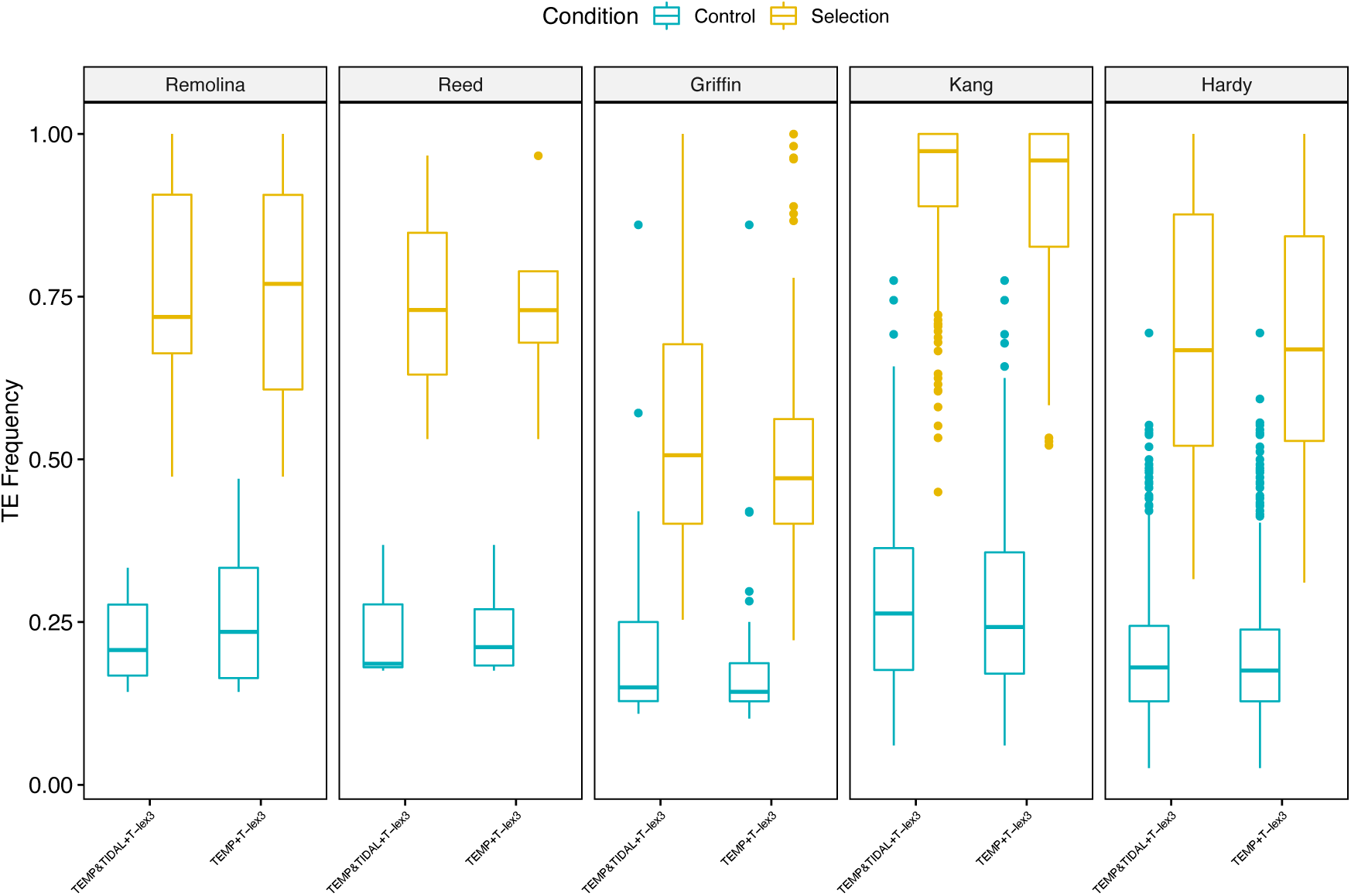
Distribution of the frequency of significant TEs in control and selection pools for each E&R study.

Using the Griffin dataset, which is the study with the highest number of replicates analyzed here, we also tested whether the frequencies of those TEs that were only identify in one selection pool (putatively absent in the mass bred, the control pools, and the other selection pools) are significantly higher than the frequencies of the TEs that were only identified in one control pool (putatively absent in the mass bred, selection pools and the other control pools). The obtained significant result (Fisher exact test: *P*-value < 2.2e-16; Figure S1) might suggest that some of the newly arisen TEs during the course of the experiment in selection pools could have been targeted by positive selection rising their frequency above those new TEs in control pools.

### Up to 35% of the candidate TEs across E&R studies are not associated with previously identified candidate SNPs or genes

Although TEs found in a single replicate (and specially if they are private *i.e.* not present in the control pools of the other replicates), and TEs detected by a single software, in the case of the non-reference TEs, were still considered as candidates to play a role in the trait of interest, they have less evidence compared with TEs present in several replicates, and non-reference TEs identified by the two software, respectively. Thus, for the rest of this work, we classified the candidate adaptive TEs in two datasets: those present in more than one replicate and detected by the two software were considered as “strong evidence TEs” –*SE*–, while the other TEs were considered as “weak evidence TEs” –*WE*–.

We first estimated the distance from the candidate TEs to the location of the closest significant SNP/gene identified in the same E&R study (see Material and Methods; Table 3). When available, we also estimated the distance from the candidate TE insertions to the location of the closest significant SNP/gene in additional E&R studies that focused on the same traits of interest. This was possible for lifespan and/or late-age fertility (Carnes, et al. 2015; Fabian, et al. 2018), and desiccation resistance (Rajpurohit, et al. 2013).

**Table 3.** Number of significant TEs with Strong Evidence (*SE*) or Weak evidence (*WE*) of selection. In Kang and Hardy, the number of significant TEs found in less than three pools are shown in parenthesis. In Griffin, for *WE* TEs, only those candidate TEs found in more than one replica or private are considered.

|  |  | Remolina | Griffin | Reed | Kang | Hardy |
| --- | --- | --- | --- | --- | --- | --- |
| **Strong evidence**<br>**(*SE*) TEs** | **Total TE number** | 0 | 10 | 0 | 37 (58) | 12 (85) |
|  | **TEs associated with**<br>**significant SNPs/genes** | 0 | 9/8 | 0 | 28 | 1 |
|  | **TEs related with trait of**<br>**interest** | 0 | 5 | 0 | 14 | 6 |
|  | **TEs in gene body region** | 0 | 70%<br>(7/10) | 0 | 73%<br>(27/37) | 83%<br>(10/12) |
| **Weak evidence**<br>**(*WE*) TEs** | **Total TE number** | 8 | 23 | 3 | 14 (78) | 4 (241) |
|  | **TEs associated with**<br>**significant SNPs/genes** | 8 | 23/21 | 0 | 9 | 2 |
|  | **TEs related with trait of**<br>**interest** | 8 | 13 | 3 | 4 | 4 |
|  | **TEs in gene body region** | 87% (7/8) | 70%<br>(16/23) | 33%<br>(1/3) | 40%<br>(6/15) | 100%<br>(4/4) |

Two of the three studies with significant TEs in the *SE* category, Griffin and Kang, showed a substantial number of significant TEs located nearby significant SNPs or significant genes (Table 3; Table S5). In Griffin, we could analyze nine out of the 10 significant TEs as the other TE was located in the fourth chromosome which was not further explored by Griffin, et al. 2017. The nine *SE* TEs were located in the same regions as significant SNPs, although not always the significant TEs and SNPs were found in the same selection pool (DR pool; Table S5B3-B4). In Kang, 28 out of 37 significant TEs are located nearby significant genes reported in Kang, et al. (2016) or in Rajpurohit, et al. (2013). The majority of these 28 TEs were in genes: i) significantly up-or down-regulated in Rajpurohit, et al. (2013); or ii) showed fixed genotypes between desiccation and control groups in Kang, et al. (2016) (Tables 3 and S5C). In addition, some of the significant TEs nearby significant SNPs or genes in the two experiments of desiccation resistance were located in the same genomic region (Table S6). On the contrary, in the Hardy study, 11 of the 12 *SE* TEs were inserted in genes different from those showing hallmarks of positive selection (Table 3 and Table S5D). Thus overall, 35% (20/58) of the *SE* TEs across studies were not associated with previously identified candidate SNPs/genes.

Regarding those TEs in the *WE* category, all significant TEs identified in the Remolina dataset were found to be associated with previously identified genes in Remolina, et al. (2012) or other studies that focused on the same trait (Tables 3 and S5A). Similarly, in Griffin, all 23 TE insertions were associated with previously identified significant SNPs (Table 3), and all but two were also associated with a significant gene (see Material and Methods; Tables S5B3-B4). On the other hand, none of the three candidate TEs identified in Reed were associated with previous significant SNPs/genes. Note that although one of the significant TEs in one of the replicas of high-sugar diet (a Transpac insertion in the 3L chromosome) was relatively close to a region with a significant SNP, this region was found significant for a different condition (high-fat diet). In Kang and Hardy, around half of the significant TEs in the *WE* category were found to be associated with previously identified significant genes (Tables 3 and S5C and S5D). Thus overall, 20% (10/52) of the significant *WE* TEs across studies were not associated with previously identified candidate SNPs/genes.

In summary, we identified both *SE* and *WE* TEs that were not linked to previously identified SNPs/genes in E&R studies (Table 3). Thus, these TEs might identify new genomic regions under selection that were overlooked when analyzing only SNPs.

### Many of the candidate TEs not associated with SNPs/genes are inserted or flanked by genes related with the traits under study

For those TEs that were not associated with previously identified significant SNPs or genes, we analyzed the function of the nearest gene to check whether these genes had been previously related to the trait of interest (Table S7). In Kang and Hardy studies, due to the high number of significant TEs, we focused on those candidate TEs found in all replicates (Table 3 and Table S7D-E). In Kang (desiccation resistance), we found five *SE* and one *WE* TEs that were within or flanked by genes showing evidence of being associated with desiccation response (Table S7D), *e.g.* the gene *α-Est10* involved in adaptation to arid environments in cactophilic Drosophila species (Rane, et al. 2019). Other genes nearby these six significant TEs were related to the insect cuticle (Pardy, et al. 2019), thermal stress (Telonis-Scott, et al. 2013; Pool, et al. 2017; Michalak, et al. 2019), salt tolerance (Davies, et al. 2014; Cohen, et al. 2020) or related with Malpighian tubule (Davies, et al. 2014), which have been all associated with desiccation response as well (Table S7D).

In Hardy (starvation resistance), we found six *SE* TEs inserted or flanked by genes related to starvation. One of the genes encompassing significant TEs, *Mics1*, is related to starvation resistance (Table 3; Table S7E) (Meng, et al. 2017). We found two genes flanking significant TEs that were also related to starvation resistance (Table 3; Table S7E): *CG1441* involved in a specific food-seeking behavior upon starvation called starvation-induced hyperactivity (Huang, et al. 2020), and *DNaseII* involved in autophagy induced by starvation (Bass, et al. 2009). Other genes (Table S6E) were involved in different diet conditions (Leow, et al. 2018) and nutritional sensing (Jayakumar, et al. 2018), olfaction behavior or olfaction memory (Arya, et al. 2015; Walkinshaw, et al. 2015) and insecticide resistance, which it has been previously shown that in some specific cases correlates with starvation resistance (Duneau, et al. 2018). We also found that the two *WE* TEs not associated with previously identified genes are associated with genes related to starvation. For instance, *cert* is involved in starvation-induced autophagy (Garlapow, et al. 2015) and *Lasp*, is a member of the diet-dependent proteins in oogenesis and it has been shown that its expression is reduced under starvation conditions (Hsu and Drummond-Barbosa 2017).

Finally, in Reed (dietary selection), we found that the three significant *WE* TEs not associated with previously identified genes are flanked by genes related to the trait of interest. In this study and in contrast to the other studies analyzed here, none of the few candidate TEs was inserted in genes with evidence of being related with nutritional status (Table 3 and Table S7B). Instead, the flanking gene of one of the candidate TEs, *CG14120*, is one of the genes that are shown to be down-regulated in response to aging and other stresses associated with sugar metabolism and proteolysis (Landis, et al. 2012). In addition, this gene is also one of those that are down-regulated as a consequence of mutations in the gene *sir2*. These down-regulated genes are enriched in pathways related to the metabolic defects such as carbohydrate metabolism (Palu and Thummel 2016). Another flanking gene, *Pld,* has a role in apoptosis induced by starvation (Maier, et al. 2019) and it has been suggested that there is a link between nutritional status and coordination of cell differentiation, a process in which *jing*, the other flanking gene of the same TE, is involved (Liu and Montell 2001). Thus overall, 17 of the 30 TEs not associated with SNPs/genes in E&R studies were located inside or nearby genes that could play a role in the trait of interest, thus suggesting that they are identifying new regions under selection overlooked by SNPs. Note that the percentage of *SE* and *WE* TEs that identify new regions under selection is similar (55% and 60%, respectively).

### Little evidence of significant TEs being enriched in specific genomic features, chromosomal arms or TE families

We explore the genomic distribution of the significant TEs and their location in terms of recombination rate and gene context. We found that many of the significant TEs are inserted in regions with relatively low recombination rates (Tables S7A-E). The number of significant TEs inserted in body regions varied from 33% to 100%, with most studies showing values above 70% (Table 3). We then analyzed whether different genomic features (*e.g.* 5’ and 3’ UTR, exon, intron, promoter and 1000 bp upstream and downstream of the CDS region), chromosomal arms or specific TE families were enriched for significant TEs. We did not find any evidence of TEs enrichment in any of the genomic features analyzed (Table S8). Except for Reed, where the number of significant TEs was small, we identified significant TEs across all chromosomal arms (Figure 2). However, only in one study, Kang, et al. (2016), we found a significant enrichment of *SE* candidate TEs in chromosomal arm 3L (Table S8D1). This is in concordance to what these authors found, with genes showing patterns consistent with hard selective sweeps preferentially located in chromosomal arm 3L. The same trend (few cases supporting strong evidence of overrepresentation) was observed for the TE family enrichment. For instance, we found that in the Kang study, the *pogo* TE family was significantly enriched in *SE* TEs when considering all significant TEs after FDR correction (Table S8D2). Likewise, in Hardy, the *FB* family is significantly enriched in the *SE* category after correcting by FDR (Table S8E3). Finally, we found weak evidence of overrepresentation for other TE families in the *WE* category and under FDR correction (Tables S8A2, S8D4).

### Candidate TEs are located nearby genes enriched for functions directly related with the trait of interest across studies

Finally, we checked whether genes located nearby all the candidate TEs identified were enriched for biological functions previously associated with the trait under selection (Falcon and Gentleman 2007). We did this considering all genes (encompassing and flanking candidate TEs) and only for genes encompassing TE insertions. While significantly enriched GO terms were found in all studies, in most of them, a higher number of significant GO terms was found when considering only genes encompassing TEs (Table S9).

In Remolina (lifespan and late-age fertility), when considering only those genes encompassing TEs, we found among others, GO terms related with oogenesis and immune response, which are key processes in modifying late-age performance at the cost of early age fecundity (Remolina, et al. 2012) (Figure 5; Table S9A2.1). These GO terms were also found in the functional enrichment analysis reported in Remolina, et al. (2012). In addition, functional terms related to antifungal activity were also found enriched in other studies on late-life fertility (Fabian, et al. 2018; Carnes, et al. 2015). In Reed (dietary selection), despite the low number of genes included in the GO enrichment analysis, and that only one of the significant TEs were within gene body regions, we found significant GO terms related to methylation involved in epigenesis (ESC/E(Z) complex; Table S9B1.3). Other significant GO terms were associated with stress granules and catalytic activity acting on RNA (Tables S9B1.3 and S9B1.2). All these terms have been associated with nutrition (Osborne and Dearden 2017) and starvation and/or malnutrition (Wang, et al. 2020; Li and Wang 2023).

**Figure 5.**
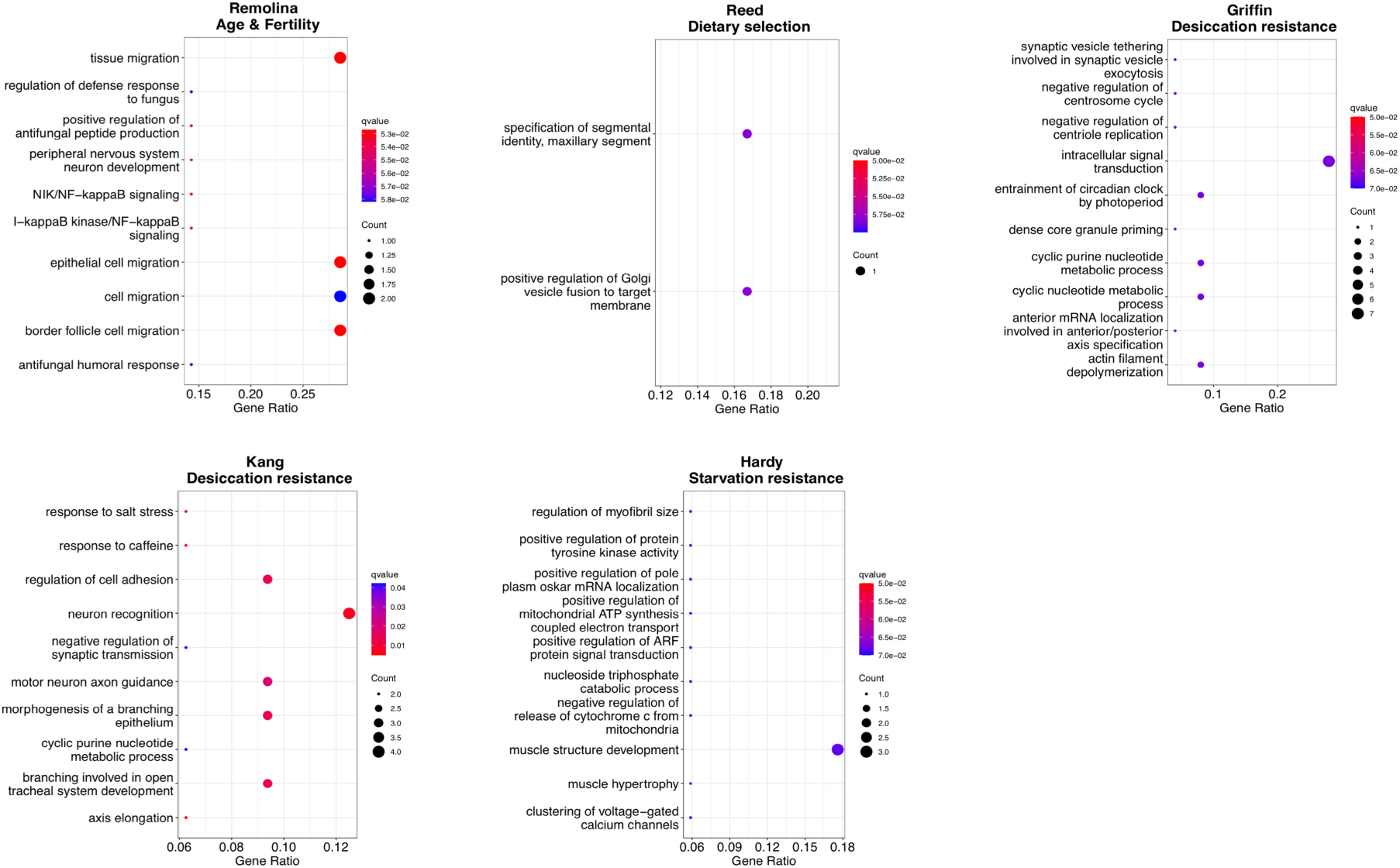
Top 10 Significantly enriched Biological Processes for each E&R studies. For all studies except Reed, the analysis was performed using the genes encompassing significant TEs

In Griffin (desiccation resistance), when the analysis was performed with genes encompassing significant TEs, we found significant enriched biological processes related with circadian clock, circadian rhythm and light stimulus, taste stimulus, ion channels and some biological processes related with calcium ion and synapsis, among others (Figure 5; Table S9C2.1). Some of the significant GO terms included not only in the biological processes category but also in the molecular function (actin binding; Table S9C2.2) and cellular component (plasma membrane; Table S9C3.2) were also previously found in Griffin, et al. 2017. This is not unexpected given that most of the significant TEs detected in our work are associated with significant SNPs and/or genes found in Griffin, et al. (2017).

In Kang (desiccation resistance), when the analysis was performed considering all genes, we only found one significant GO term (Table S9D1.1). However, when the analysis was performed considering only the genes encompassing TEs, we found some significant GO term similar to those found in the other desiccation study analyzed (Griffin, et al. 2017), such as those related to the visual system, synapsis, and chemical stimulus related with taste (Table S9D2.1). Besides, we found some relevant GO terms not found in either of the two desiccation studies analyzed in this work, such as those involved in tracheal system development, and response to salt stress (Figure 5; Table S9D2.1).

Finally, in Hardy (starvation resistance), we did not find any significant GO term when the analysis was performed considering all genes. However, when we performed a GO analysis considering genes encompassing TEs, we found significant terms related with muscle development and morphology and biological processes involving mitochondrial function (Figure 5; Table S9E2.1). There is a well-documented body of evidence describing the interplay between starvation and longevity, since dietary restriction extends lifespan in multiple species (Longo and Anderson 2022). However, in Drosophila, it has also been documented an interplay between longevity, starvation and muscle function. For instance, it is possible to delay systemic aging via preservation of proteostasis and muscle function (Demontis and Perrimon 2010) and also, there is a muscle-specific mutant in *Drosophila* which exhibits resistance to starvation (Guo, et al. 2023). In addition to the effect on muscle function, dietary restriction can also modify the mitochondrial activity and morphology through changes in the expression of genes involved in electron transport chain and enzymatic activity (Zid, et al. 2009). Overall, genes located nearby candidate TEs are enriched for GO terms associated with the trait of interest. While some of these GO terms were already identified based only on the analysis of SNPs, the analysis of TE insertions allowed to identify new GO terms likely involved in adaptation to the selective condition.

## DISCUSSION

In this work, we searched for evidence of TEs driving the response to different selection regimes in five different E&R studies (Table 1). We focused on those TEs that significantly increase their frequency when selected for the trait of interest over the time of the experiment. Our results showed that TEs are candidate selection targets in response to the four selective regimes analyzed (Table 2). Some of the identified candidate TEs are located nearby previously identified candidate SNPs (Table 3). In this case, disentangling which variant is the true causative loci responding to selection is not trivial. However, we argue that because TEs are known to generate mutations of large effect, which are likely to be deleterious, some of those TEs that significantly increase in frequency after a few generations of selection are likely to be the causal mutation. On the other hand, 35% of the identified TEs are not associated with previously identified candidate SNPs (Table 3). Note that, although E&R experimental populations often show long-range blocks of linkage disequilibrium, the distance between candidate TEs identified and the previously identified SNPs was greater than the linkage disequilibrium blocks reported (1 Mb to 2.6 Mb depending on the data set; see Material and Methods). The majority of TEs that are not linked to previously known significant SNPs (17/30) are located nearby genes involved in functions that have been previously associated with the trait of interest (Table S5). This suggests that the selected regions only identified by TEs are likely to play a role in the evolution of these traits. Thus, analyzing TEs in E&R studies leads to the identification of regions under selection that went undetected when analyzing only SNP variants, with some of these regions identifying genes not previously related to the trait of interest. As such, including TEs in E&R studies should provide a more complete picture of the genetic basis of adaptation.

One of the objectives of our analysis was to assess if TE insertions show parallel patterns of selection across replicates as has been shown for SNPs. Identifying patterns of parallel selection allows evaluating how predictable adaptation is (Schlotterer 2023). Indeed, we found that for three of the five studies analyzed, the level of replicability of TEs was similar or higher than that of SNPs (Figure 3 and Table 2). We further showed that the level of replicability observed is likely to be underestimated as a substantial proportion of the TEs only found in one of the selective replicates was not present in the control replicates most probably due to the stochasticity in the sampling of individuals from the starting population (Figure 3 and Table 2).

Taking into account the genes located nearby all candidate TEs identified, those linked to previously identified SNPs and those identifying new regions under selection, we found that they are enriched for functions that have previously been related with the trait under study, further suggesting that these insertions are bona fide selection targets. In some cases, *e.g.* Remolina, et al. (2020), some of the genes associated with the candidate TE insertions and candidate SNPs, being the latter identified in the same or other studies analyzing the same trait, were enriched for the same GO terms (Fabian, et al. 2018; Carnes, et al. 2015). In other studies, we identified genes enriched for GO terms that were not initially identified by SNPs of the same study but these same GO terms were associated with the trait of interest using other experimental approaches. This is the case of genes related to the tracheal system development identified in Kang, et al. (2016) that were previously identified using a QTL approach, artificial selection approaches, and analysis of mutant strains (Bradley, et al. 1999; Gomez, et al. 2015; Wang, et al. 2018). Thus, incorporating the analysis of TEs into E&R studies also gives a more complete picture of the functions associated with the selective regime under study.

Because we analyzed two E&R desiccation resistance studies, we were also able to look for instances of different TE insertions targeting the same genes as has been previously described (Niu, et al. 2019). Note that populations used in these two studies were collected from distant geographical locations: Jabalpur, India (Kang, et al. 2016) and Melbourne, Australia (Griffin, et al. 2017). We found evidence for candidate TE insertions, located in different gene regions and belonging to different TE families, targeting the same gene across studies (Table S7). We cannot discard that TEs targeting the same genes were due to some particular characteristic of these genomic regions that made them more prone to TE insertions. However, we argue that these TEs are good candidates for follow up functional analysis as they might be inducing similar effects on the nearby gene.

Although we focused on those available E&R studies with more power to detect the causal selected mutations, our analysis is likely underestimating the role of TEs in the traits analyzed. First, there is evidence showing that the identification of TEs based on short-read sequencing fails to identify up to 57% of the insertions present in a genome (Rech, et al. 2022). Additionally, we are only analyzing those TEs that are present at 10% frequency in the populations, as this is the threshold suggested in Nelson, et al. 2017 for TEMP, one of the software used to identify *de novo* TE insertions, to avoid false positive sequences. Thus, we are only considering a subset of all the TEs present in the populations analyzed. Second, for those studies where the ancestral population is available (mass bred/baseline pool), we detected a higher number of TEs in control/selection pools than in the mass bred or baseline pools, which may suggest that some TEs are mobilized, and thus generate *de novo* mutations, during the course of the experiment. Indeed, for the Griffin dataset (Griffin, et al. 2017), we found a significant increase in frequency of those TEs that have potentially moved in the selection pools (Figure S1). This result suggests that it might be worth analyzing those TEs that are mobilized during the course of the experiment as potential candidates to be targets for selection. Finally, TEs are mostly present at low population frequencies, which makes it more challenging to detect significant increases in frequencies in response to selection in comparison to SNPs that are present at higher frequencies (Brennan, et al. 2019). Thus, to improve our capacity of detecting TEs as targets of selection in E&R experiments, we should take into account the idiosyncrasy of TE variants: difficult to detect based on short-read sequencing, ability to move and generate new mutations, present at lower frequencies compared with SNPs. Given the continuous decrease in long-read sequencing prices and the improvement of the DNA extraction protocols (*e.g.* Kim, et al. 2023), using a combination of short-and long-read techniques to sequence E&R experiments will increase our ability to detect TE insertions as targets of selection.

Increasing population sizes is recommended to maximize the power to detect selected variants (Kofler and Schlötterer 2014), and it is particularly relevant for TE insertions as most of these variants are present at low population frequencies. Besides increasing population sizes, increasing the number of replicate populations would also be particularly relevant for the analysis of TE insertions as it will allow the detection of newly mobilized insertions. However, determining the presence/absence of candidate TEs in the mass bred or baseline would eventually be necessary to discern between those TEs not detected in some of the pools because of their low frequency or because they are newly arisen, *e.g.* by using a PCR approach if the number of candidates is not too big. Finally, candidate TEs could then be tested by deleting the selected TEs using genome editing techniques such as CRISPR/Cas systems (Merenciano, et al. 2023) and performing experimental assays to verify their adaptive nature (*e.g.* Ullastres et al 2021, Merenciano and González 2023).

Overall, in this study, we identify TEs that are bona fide targets of selection in E&R studies analyzing different traits: stress response, life history and dietary selection. Given that TEs are a valuable source of variation and their proven implication in rapid adaptation (Stapley, et al. 2015; Baduel and Quadrana 2021), we argue that, beside SNPs, TEs should also be considered in E&R studies.

## MATERIAL AND METHODS

### *D. melanogaster* sequenced data

We downloaded the FASTQ files corresponding to the *Drosophila melanogaster* Pool-seq data generated in five different E&R experiments where the genetic basis of adaptation to selection for increased lifespan and late-age fertility (Remolina, et al. 2012), different diets (Reed, et al. 2014), desiccation (Kang, et al. 2016; Griffin, et al. 2017), and starvation resistance (Hardy, et al. 2018) were studied. The Pool-seq data corresponds to mass bred and baseline populations and to different numbers of replicates of control and selected populations (Table 1).

### Mapping

For each pool, we filtered the raw FASTQ reads to remove low-quality bases with minimum base PHRED quality = 18 using *cutadapt* v. 1.8.3 (Martin 2011). Trimmed reads were mapped against the *D. melanogaster* reference genome *v.* 6.12 and to other sequences from common commensals and pathogens as in Kapun et al. 2020 using *BWA* (mem algorithm) v. 0.7.16a-r1181 (Li and Durbin 2009). We used *Picard-tools* v. 2.8.3 (http://broadinstitute.github.io/picard) to remove duplicate reads, and *GATK* v. 3.7-0-gcfedb67 (McKenna, et al. 2010) to re-aligned sequences flanking insertions-deletions (indels). Finally, we filtered reads due to a possible *D. simulans* contamination as in Kapun et al. 2020. *SAMtools* v. 1.6 (Li, et al. 2009) was used to retrieve autosomal and sex chromosomes information and to generate *pileup* or *mpileup* outputs from BAM files.

### Reference and non-reference TE insertions identification

We used T-lex3 (Bogaerts-Márquez, et al. 2020) to detect and to obtain the frequencies of TE reference insertions. We ran T-lex3 using default parameters with -pairends no and adjusting the -A parameter for the specific read length of each data set. In the case of the Reed data set, T-lex3 did not work using default parameters so we used less stringent parameters: -minQ 20 and -id 80. However, we obtained similar results, so the reference TEs could not be analyzed for this specific dataset. For TE frequency estimation we used the default -freq, -pooled parameters and we adjusted -maxR using the 0.9 quantile of sample coverage.

We used two different software, *TEMP* v.1.05 (Zhuang, et al. 2014) and *TIDAL* v.1.0 (Rahman, et al. 2015), that use short-read sequencing to detect non-reference insertions (present in Pool-seq data samples and absent in the reference genome). *TEMP* uses the information provided by both pair-end and split reads to infer the TE insertion interval at nucleotide resolution whereas *TIDAL* uses single-end and split reads (*TIDAL* automatically converts all pair-end into single-end reads). The *TEMP* module insertion was run with command line options -m 3 -c 8 -x 30 for all datasets and adjusting -f parameter for the specific insert size of each data set. *TEMP* uses a TE consensus fasta file and a bed file containing the genomic coordinates of all annotated TEs. These two files were downloaded from Repbase and from UCSC Genome Browser respectively, as indicated by the authors (Zhuang, et al. 2014). *TIDAL* was run using the same annotation files and parameters as in the default settings (Rahman, et al. 2015). The - min_len parameter was adjusted to the specific insert size of each data set.

To avoid including false positive TE insertions, we only considered as reliable those insertions with split-read support at both ends of the insertion (“1p1”) and with sample insertion frequencies above 0.10 for *TEMP* (Zhuang, et al. 2014; Nelson, et al. 2017). All predictions made by *TIDAL* were considered as reliable since potentially false positives were filtered out by using BLAT score with default parameters. We used a dictionary to standardize the name of the TE families since the two software uses different nomenclature for TE family names (Table S10).

We used the R packages *GenomicRanges* v.1.34.0 (Lawrence, et al. 2013), *tidyr* v.0.8.3 (Wickham and Henry 2020) and *dplyr* v1.1.2 (Wickham et al. 2023) to find those TE insertions detected by the two software. We considered as the same TE insertions those in which the genomic intervals predicted by both programs: i) are in the same chromosomal arm; ii) their start/end coordinates overlaps; iii) the predicted insertion corresponded to the same TE family.

### Detection of candidate TEs to be under positive selection

We searched for those TEs that significantly increase in frequency from control to selection pools or from mass bred or baseline to selection pools, and among those, we identified the significant TEs that were shared by selection replicates. We also checked the function of the genes where the TE is inserted or the closest genes to the significant TE insertions (Flybase: Gramates et al. 2022), the recombination rate where the significant TEs were inserted and their distance from significant SNPs/genes described previously in each E&R experiment. Similar to some E&R experiments that used SNPs, the significance of the change in frequency was determined using a Fisher’s exact test and applying a Bonferroni cutoff of 0.05. We used this approach since it was previously shown that the performance of this test was the most conservative compared with other conditions such as likelihood ratio test (Lynch, et al. 2014) or the allele frequency change expected due to genetic drift alone (Griffin, et al. 2017). Fisher’s exact test was computed using the function *fisher.test* from package *stats* v. 3.5.2. Non-reference TEs were analyzed separately according to whether they were found by both (TIDAL and TEMP) or only one software (TEMP), since TEMP is the only software used here that estimates the TE frequencies.

The strategy to find candidate TEs was slightly different for some datasets because the design of the different E&R experiments from which the data were generated was also different: Remolina, et al. (2012): the six Pool-seq data corresponds to three paired replicates of control-selection *D. melanogaster* populations. From those TEs detected in each pool (see above), we search for those TEs that fulfill two conditions: i) were present in each paired control-selection replicate; and ii) increase significantly in frequency between the control and the selection pool. Finally, we identified those candidate adaptive TEs that were shared by more than one selection replicate.

Reed, et al. (2014): the 13 Pool-seq data corresponds to seven pools of a baseline *D. melanogaster* population and two replicates of three evolved populations under different diets. From those TEs detected in each pool, we search for those TEs that fulfill two conditions: i) were present in the baseline population and each replicate of the three evolved populations; ii) change significantly in frequency between baseline population and each replicate of the three evolved populations. Finally, we identified those TEs that were shared between replicates/populations of each evolved population.

Griffin, et al. (2017): the 11 Pool-seq datasets correspond to one mass bred and 5 replicates of control-selection *D. melanogaster* populations. As in Griffin, et al. (2017), to generate the single mass bred population, we down sampled four independently sequenced pools in order to have the same number of total reads as in the selection/control pools. First, from those TEs in each pool, we search for those TEs that fulfill the following conditions: i) were present in the mass bred pool and each control and selection replicate; ii) change significantly in frequency between mass bred and each control pools (*P*-value < 0.05; no multiple testing correction) and mass bread and selection pools (Bonferroni cutoff of 0.05). Then, also as in Griffin, et al. 2017, we exclude those TEs that change significantly in frequency between the mass bred and the control pools from the list of TEs that significantly change in frequency between the mass bred and selection pools since those TEs are considered to be involved in adaptation to the laboratory conditions. Finally, we identified those candidate adaptive TEs that were shared by more than one selection replicate.

Kang, et al. (2016) and Hardy, et al. (2018): the six Pool-seq data in each experiment corresponds to three replicates of control-selection *D. melanogaster* populations. For each experiment, from those TEs detected in each pool, we search for those TEs that fulfill two conditions: i) were present in control pools and each selection replicate; ii) change significantly in frequency between controls and each selection replicate. Finally, we identified those TEs that shared by more than one selection replicate.

For all the data sets analyzed, we grouped the TEs that significantly increased in frequency between control and selected conditions according to its degree of supporting evidence: those found in more than one replica were considered to have more support to be under positive selection (TEs with strong evidence –*SE*–), and those found in a single replica were considered to have less support (TEs with weak evidence –*WE*–). For non-reference TEs, besides the number of replicates in which they were present, we also required that the TE was identified by the two softwares to be considered a TE with strong evidence. In the case of Remolina and given that the control-selection replicates are paired, those TEs showing a significant increase in frequency in one pool but a non-significant increase in the rest of replicates were not considered in the analysis.

Candidate TEs were then classified as associated and non-associated to significant SNPs or genes previously reported in the same or other related studies analyzing the same trait. A candidate TE is considered to be associated to significant SNPs or genes if the distance between the coordinates of the candidate TE and the closest significant SNP/gene previously reported is less than the length of the haplotype blocks reported in the same or other related studies focused on the same trait (lengths up to: 1.8 Mb in Remolina, et al. 2012; 2.6 Mb in Carnes, et al. 2015; 1 Mb in Griffin, et al. 2017; 2 Mb in Kang, et al. 2016 and in Hardy, et al. 2018). In Griffin, because this study has a relatively large number of replicates compared to the other studies, we only considered as candidates those TEs found significant in more than one replicate and those that are private. Similarly, in Kang and Hardy, due to the elevated number of candidate TEs found, we only considered those candidate TEs present in all pools.

### Enrichment analysis: chromosome, genomic features and TE families

We tested if specific chromosomes, genomic features or TE families were enriched with significant TEs. We used the R package *GenomicFeatures* v. 1.34.3 (Lawrence, et al. 2013) and DescTools: 0.99.41 (Signorell et mult. al. 2021) to update the GFT file for *D. melanogaster* reference genome v6.35, creating features for introns, 1-kb flanking, 5’UTR, and 3’UTR regions and use this updated GTF to retrieve the features of the significant and non-significant TEs. Fisher test using the function *fisher.test* from R package *stats* v. 3.5.2. was computed to test for the enrichment in significant TEs of specific chromosomes, genomic features or TE families. The enrichment analysis was performed grouping the significant TEs according to the *SE*/*WE* classification. For the TE family enrichment analysis, only TE families with more than 20 copies in the genome were considered. The recombination rate where the significant TEs were inserted was calculated using the recombination rate calculator (RRC; Fiston, et al. 2010) available at http://petrov.stanford.edu/cgi-bin/recombination-rates_updateR5.pl. We first used the genomic tool *Coordinates Converter* in Flybase (https://flybase.org/convert/coordinates) to convert the TE insertion coordinates of release 6 of D. melanogaster reference genome to those corresponding to release 5 since the RRC is based on release 5.

### GO analysis

We used the R packages *GSEABase* v.1.60.0 (Morgan, et al. 2022), *GOstats v.2.64.0* (Ashburner et al 2020; The Gene Ontology Consortium 2023; Falcon, et al. 2007) and *org.Dm.eg.db* v.3.16.0 (Carlson 2019) to perform the GO enrichment analysis. The obtained P-values were transformed into q-values using the R package *q-value* v.2.30.0 (Storey, et al. 2022) to correct for multiple testing. The cutoffs for considering a GO term significant was set to 0.01 and 0.10 for the P-values and the q-values respectively. GO enrichment analysis was performed using two different set of genes for each experimental data: i) the genes where the candidate TEs are inserted or both the flanking 5’ and 3’ genes (in case that the candidate TE is inserted in intergenic region); ii) only those genes with inserted candidate TEs.

## ACKNOWLEDMENTS

We thank members of the González lab for providing comments on a previous version of this manuscript. J.G. is funded by grant PID2020-115874GB-I00 funded by MICIU/AEI/10.13039/501100011033 and by grant 2021 SGR 00417 funded by Departament de Recerca i Universitats, Generalitat de Catalunya.

## Data availability

All the sequences used in this study were downloaded from public databases (NCBI). The BioProject accessions are: PRJNA185744 (Remolina), PRJNA194129 (Reed), PRJNA306702 (Griffin), PRJNA304655 (Kang), PRJNA315172 (Hardy).

## Code availability

The scripts used in this study have been deposited to GitHub and are freely accessible from: https://github.com/GonzalezLab/TEs_Evolve-and-Resequencing

